# Variation in upstream open reading frames contributes to allelic diversity in protein abundance

**DOI:** 10.1101/2021.05.25.445499

**Authors:** Joseph L. Gage, Sujina Mali, Fionn McLoughlin, Merritt Khaipho-Burch, Brandon Monier, Julia Bailey-Serres, Richard D. Vierstra, Edward S. Buckler

## Abstract

The 5’ untranslated region (UTR) sequence of eukaryotic mRNAs may contain upstream open reading frames (uORFs), which can regulate translation of the main open reading frame (mORF). The current model of translational regulation by uORFs posits that when a ribosome scans an mRNA and encounters a uORF, translation of that uORF can prevent ribosomes from reaching the mORF and cause decreased mORF translation. In this study, we first observed that rare variants in the 5’ UTR dysregulate protein abundance. Upon further investigation, we found that rare variants near the start codon of uORFs can repress or derepress mORF translation, causing allelic changes in protein abundance. This finding holds for common variants as well, and common variants that modify uORF start codons also contribute disproportionately to metabolic and whole-plant phenotypes, suggesting that translational regulation by uORFs serves an adaptive function. These results provide evidence for the mechanisms by which natural sequence variation modulates gene expression, and ultimately, phenotype.

## Introduction

Understanding the mechanisms by which genetic variation produces phenotype will depend on learning how gene expression is regulated differently between alleles. Although moderate correlations between mRNA and protein have been observed across genes (r =0.3-0.8; reviewed in(1)), the correlation across individuals tends to be quite low (r< 0.25)(2, 3). Additionally, there is only modest overlap between variants associated with mRNA levels and variants associated with protein levels (2, 4–6). Together, these findings suggest an extensive role of post-transcriptional regulation in determining gene expression. Protein synthesis is energetically expensive, which implies high selective pressure on individuals to produce the appropriate quantity of each protein(7, 8). Despite this strong selective pressure, protein levels are heritable and vary between individuals(5, 9), which means that there must be adaptive advantages to allelic differences in protein abundance. Motivated by these observations, an outstanding question is: how does genetic variation contribute to variation in protein abundance?

Secondary structure and high GC content of the 5’ UTR can affect translation efficiency, presumably by the formation of secondary structure that obstructs scanning of the 43S pre-initiation complex and completion of initiation(10–13), a mechanism which can regulate gene expression response to environmental conditions(14). However, single base-pair substitutions (the focus of this study) do not cause appreciable changes in secondary structure or GC content of the 5’ UTR. Upstream open reading frames (uORFs) located in the 5’ UTR and preceding the main open reading frame (mORF) also contribute to post-transcriptional gene regulation, generally by reducing translation of the mORF(12, 13, 15). Studies in human have shown that variants which create or disrupt uORFs are under negative selection and associated with disease phenotypes (16, 17). Across eukaryotes, the strength of uORF Kozak sequence is predictive of translation initiation efficiency (18). While the effects of uORF translation on protein abundance have been demonstrated in several genes (15, 16), or across alleles in massively parallel reporter assays (12, 13), less is known about the genomewide effects that natural sequence variation in uORFs has on protein abundance.

Previous studies of genetic variability for protein abundance between individuals(2, 4, 6, 19, 20) have primarily used genomewide association studies (GWAS), an approach that is most effective for alleles that are at high frequency in the population being studied. To be sufficiently powerful, GWAS require large population sizes and are still generally best suited to identifying common alleles, which tend to have smaller effect sizes but can be important for locally adapted phenotypes. In contrast, rare variants tend to have larger, deleterious effects(21–23) but are difficult to identify by GWAS. Because rare alleles 1) are more likely to have large effects, and 2) have been implicated in dysregulation of mRNA abundance(24, 25), we reasoned that in aggregate, rare alleles would be ideal candidates for identifying specific sets of variants that contribute most to post-transcriptional regulation. Our findings for rare alleles could then be validated in the pool of common alleles.

In this study, we answer the preceding questions with maize (*Zea mays* L.), which has large homogeneous tissues that facilitate high quality proteome extraction; extremely high levels of phenotypic and genetic diversity(26, 27); and rapid linkage disequilibrium decay that allows high genetic resolution with fewer individuals(28, 29). We use two distinct sets of diverse inbred maize lines: a technically and biologically replicated set of four inbreds, and a mostly unreplicated set of 95 inbreds previously described(2). We first use rare alleles, which generally have larger and more disruptive effects, to learn that variants in the 5’ UTR, specifically those which modify uORFs, have the most influence on protein abundance. We then use those findings to identify and test common variants which likely have weaker but adaptive effects, and demonstrate their effects on protein abundance as well as their contribution to metabolic and whole-plant phenotypes.

## Results & Discussion

Regulation of gene expression is largely genetically determined; mRNA abundance is heritable in organisms across kingdoms(30), as are proteomic(5, 9) and metabolite levels(31). Quantification of 7,524 peptides in a pair of leaves in developmentally matched juvenile plants (four replicated maize inbreds; Supplemental Data 1) confirmed that protein abundance is heritable in maize, with median heritability of 0.79, similar to mRNA and metabolite abundances(31) (Supplemental Figure 1). These high heritabilities not only demonstrate low measurement error, but also reveal extant adaptive genetic variation in protein abundances.

We identified rare variants based on a minor allele frequency (MAF) < 0.02 in the maize HapMap3.2.1 population, which contains >1,200 varieties of maize and its wild relatives from around the world(32). Rare variants were classified based on their location within five genic features: promoter, 5’ UTR, coding sequence (CDS), intron, or 3’ UTR (Figure 1b). In each genic feature, we tested for differing protein abundance among 95 diverse inbred lines (Supplemental Data 2), classified at each SNP by whether they have the common or rare allele. Introns act as in control, as they should be unable to exert post-transcriptional regulation due to their typical absence from the mRNA; we observe a corresponding lack of association between intronic rare alleles and protein abundance (Figure 1a). Rare alleles in the promoter and 5’ UTR were associated with more variable protein abundance, with rare alleles in the 5’ UTR showing the strongest effect (Figure 1a,c, Supplemental Figure 2). This finding reinforces previous implication of the 5’ UTR in post-transcriptional regulation(10, 12, 13, 16–18, 33, 34), and the lack of any significant effect from variants in the 3’ UTR contrasts with recent work focused on the role of the 3’ UTR in translational regulation in maize(35).

**Figure 1:**
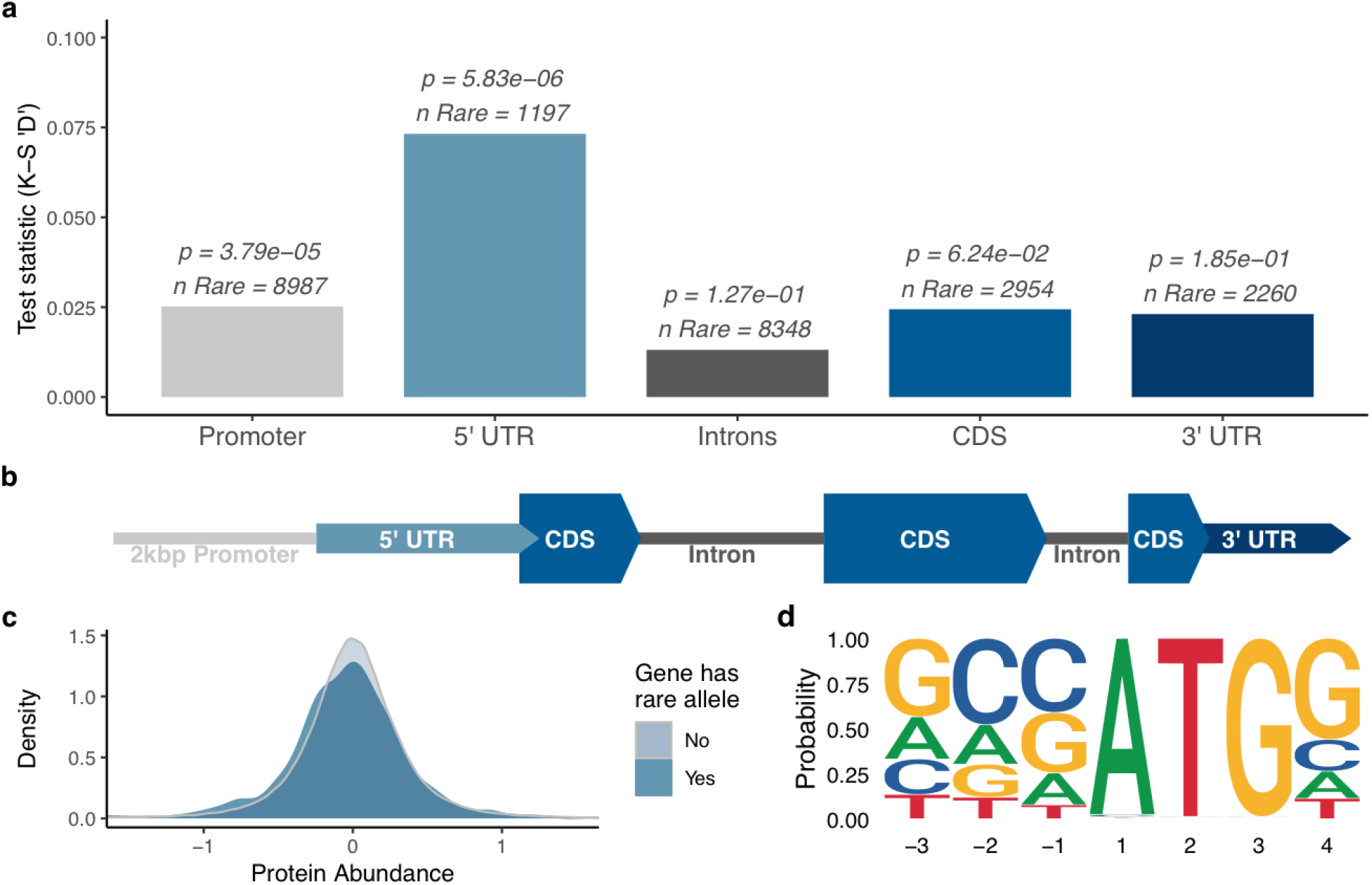
Individuals with rare alleles have more variable protein abundance. (a) Individuals with at least one rare allele (MAF < 0.02) in each genic region (b; generic gene) were compared to individuals with no rare alleles in any of the genic regions. The Kolmogorov-Smirnov (K-S) test statistic ‘D’ indicates differences in the distribution of protein abundance between individuals with and without rare alleles. Rare alleles in the 5’ UTR are significantly associated with dysregulated protein abundance (two sided K-S test, number of observations without rare allele = 184,668). (c) Distribution of protein abundance for individuals with rare alleles in the 5’ UTR (blue), compared to individuals without rare alleles (light blue). (d) Empirical maize Kozak sequence derived from the predicted start codon region of annotated gene models in v5 of the B73 genome.

The 5’ UTR can contribute to post-transcriptional regulation of gene expression, and machine learning models have been successful at predicting protein abundance from sequence. However, application of previously developed machine learning models(12, 36) proved unable to predict allelic differences in protein abundance from 5’ UTR and transcript sequence in four maize inbreds. Instead, we turned to evaluating mechanisms by which these observed rare variants might be disrupting translation.

We hypothesized that rare alleles can disrupt existing uORF start codons or create start codons that generate novel uORFs, thereby decreasing or increasing (respectively) uORF translation and ultimately altering translation of the mORF (Figure 2a,b). Based on this hypothesis, and under the assumption that there is an inverse relationship between uORF and mORF translation, we predicted that 1) rare alleles that weaken existing uORF start codons will be associated with greater mORF protein abundance (Figure 2a), and 2) rare alleles that cause new or strengthen existing uORF start codons will be associated with lower mORF protein abundance (Figure 2b).

**Figure 2:**
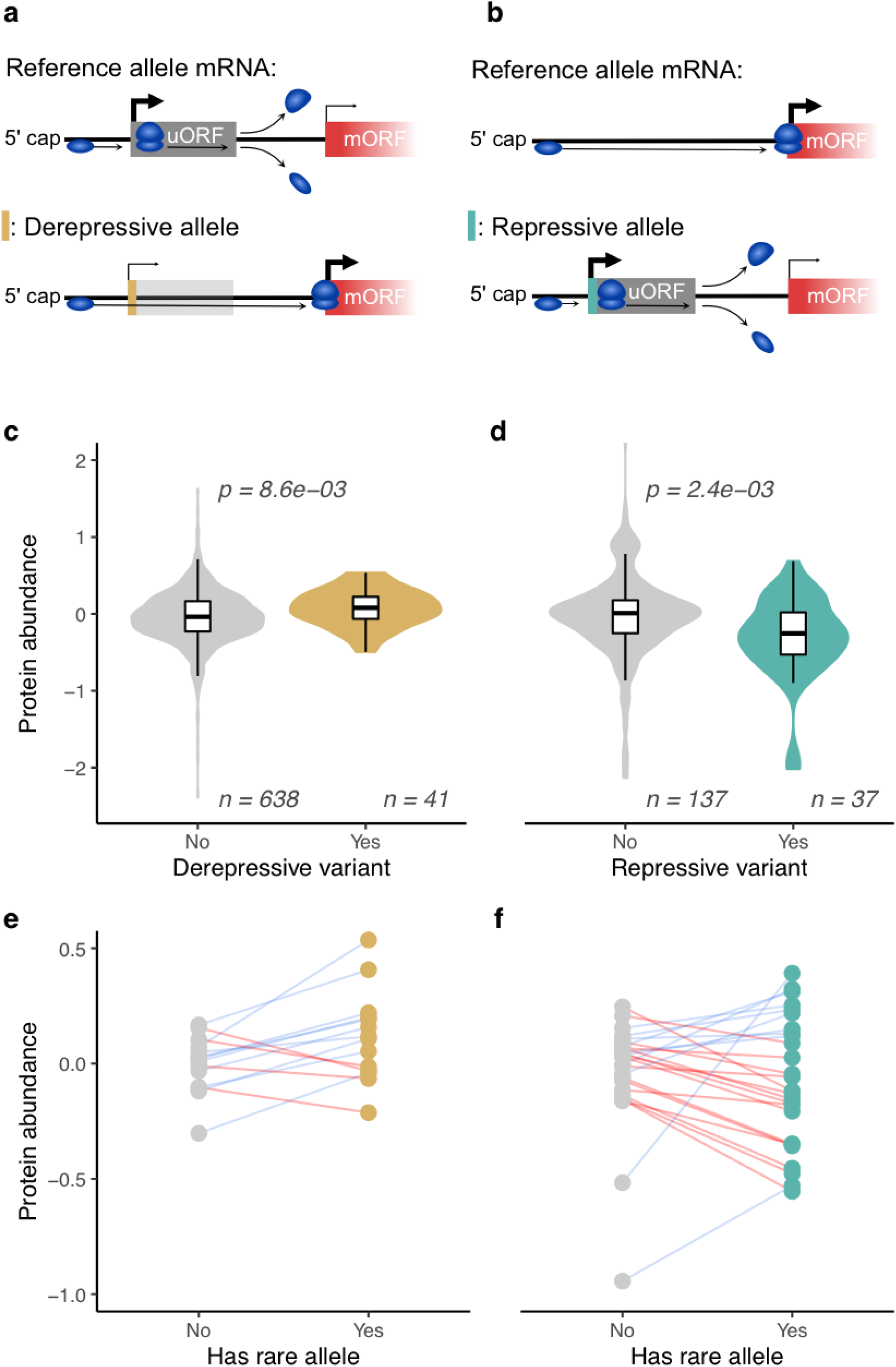
Rare variants that alter upstream open reading frame start codons are associated with altered protein abundance of the main open reading frame. (a) Rare SNP alleles which disrupt or weaken the start codon of an existing translated upstream open reading frame (uORF) are associated with derepression of the main open reading frame (mORF), whereas (b) rare alleles which cause a new start codon or strengthen an existing start codon in the 5’ UTR are associated with repression of the mORF. c) and d) show the effects of derepressive and repressive rare alleles, compared to the effects of other rare alleles in similar contexts (two sided Mann-Whitney test; boxplots show median, first and third quartiles, and whiskers extend no further than 1.5 times the interquartile range). e) and f) compare protein abundance between individuals with rare and common alleles on a per-gene basis. Individuals with rare alleles of derepressive variants often have greater protein abundance than individuals with the common allele. Individuals with rare alleles at repressive variants often have lower protein abundance than individuals with the common allele. Each pair of points connected by a line represents a single gene; the points are the median protein abundance for individuals with either the rare or common SNP allele; connecting lines are colored by whether the rare allele increases (blue) or decreases (red) protein abundance.

To test the first prediction, we identified uORFs that show evidence of translation based on ribosome profiling data(37), then searched for rare alleles within or near the start codons of those uORFs. A score was assigned to the common and rare alleles based on similarity of their surrounding sequence to the maize Kozak sequence (Figure 1d)(38), and variants with rare alleles that weaken the Kozak sequence were labelled as ‘derepressive variants’, based on hypothesized effect of the rare allele on mORF protein abundance (Supplemental Data 3). Derepressive variants were compared to rare alleles anywhere else in the 5’ UTRs of genes with translated uORFs, including within uORFs themselves but excluding the start codons of translated uORFs. This comparison revealed that derepressive variants are associated with 31% greater mORF protein abundance than rare alleles elsewhere in the 5’ UTR of genes with translated uORFs (Figure 2c; p=8.6e-03, two-sided Mann-Whitney test). We also studied the difference in protein abundance between individuals with the common and rare derepressive alleles. In 10 out of 14 cases, the rare derepressive alleles were associated with increased mORF protein abundance relative to the common allele (Figure 2e).

To test the second prediction, we performed similar analyses to those described above but focused on genes with either no annotated uORFs, or with annotated uORFs that did not show any evidence of translation(37). We searched for rare alleles that increased their surrounding sequence’s similarity to the maize Kozak sequence(38), and labelled them as ‘repressive variants’, based on their hypothesized effect on mORF protein abundance (Supplemental Data 3). We compared repressive variants to rare alleles which are located in genes with no annotated or translated uORFs, but decrease the surrounding sequence’s similarity to the Kozak motif. Repressive variants are associated with a 45% decrease in protein abundance of the corresponding mORF (Figure 2d; p=2.4e-03, two-sided Mann-Whitney test). Comparing individuals with rare alleles at repressive variants to individuals with common alleles revealed 17 out of 26 rare alleles were associated with decreased mORF protein abundance (Figure 2f).

Given that rare, putatively deleterious variants appear to dysregulate mORF protein abundance by altering start codons of uORFs, we wondered whether historical mutations with similar mechanisms may have conferred adaptive advantage and risen to a higher allele frequency via positive selection. We identified common (MAF > 0.1) repressive and derepressive variants using the same criteria described above (Supplemental Data 4), and tested their association with mORF protein abundance. We found an enrichment of significant associations between repressive or derepressive variants and mORF protein abundance (Figure 3a; one-sided Mann-Whitney test). We reasoned that if these common alleles are adaptive, they may also affect metabolic or physical phenotypes. Indeed, derepressive variants show greater than 16% enrichment for GWAS hits over all common variants in 5’ UTRs, with > 77% of derepressive variants associated with at least one phenotype. The enrichment of GWAS hits among derepressive variants may explain previous observations that the 5’ UTR has an outsized contribution to quantitative traits in maize(39). On the other hand, repressive variants show a statistically insignificant 5% enrichment (Figure 3b; two-sided binomial test; Supplemental Data 5). The enrichment of GWAS hits for metabolic phenotypes is particularly notable because uORFs have been implicated in metabolite based regulation involving the translating ribosome(40, 41). Derepressive variants show stronger association with protein abundance and greater enrichment for GWAS hits than repressive variants, possibly because reduced translation of the mORF may be under strong negative selection(42), or because SNPs are more likely to disrupt existing uORFs than to create new translated uORFs(43).

**Figure 3:**
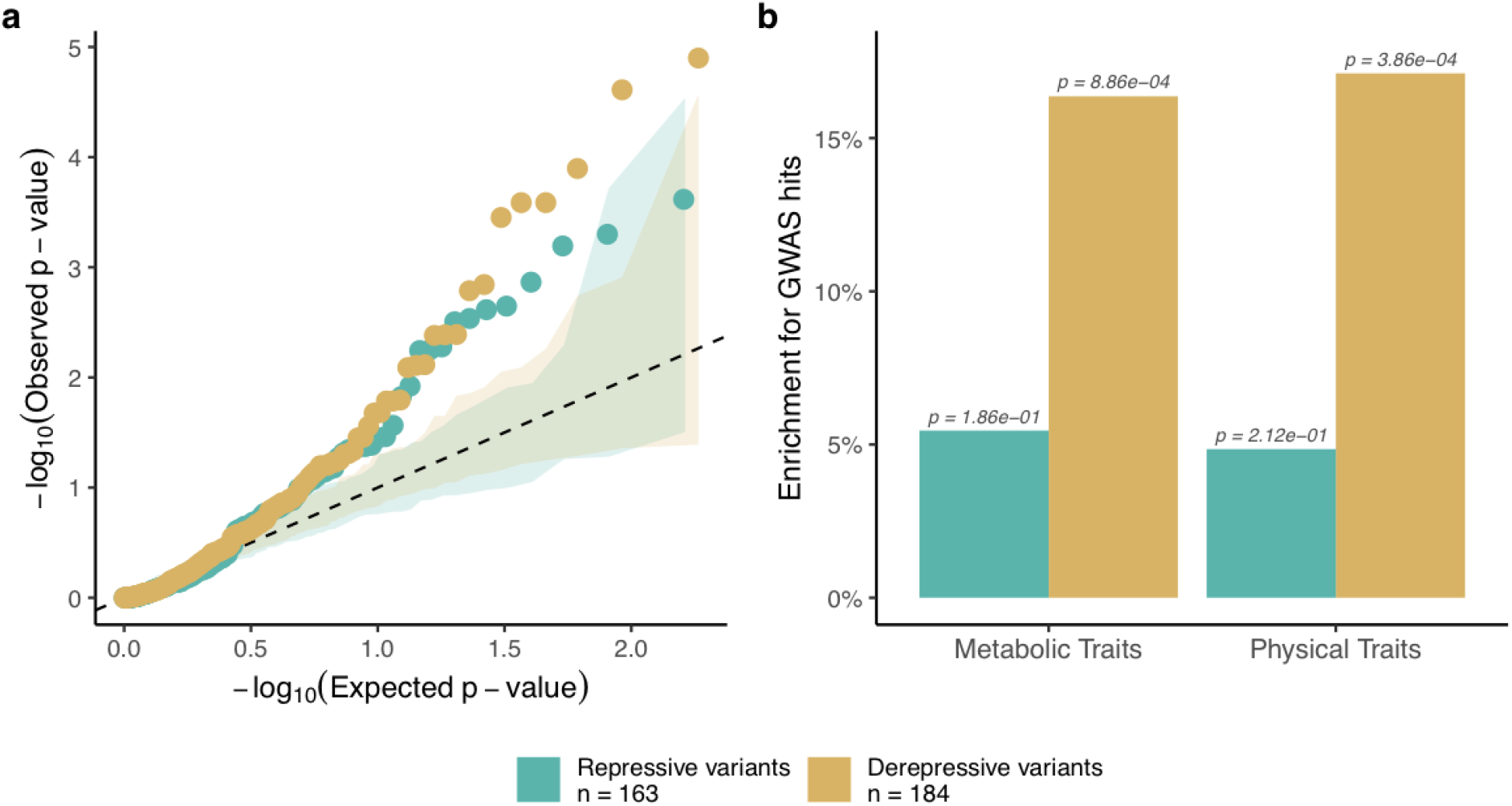
Common variants that alter upstream open reading frame start codons show evidence of adaptive function. a) Common variants (MAF > 0.1) show significant association with main open reading frame (mORF) protein abundance based on one-sided Mann-Whitney test for increased (derepressive variants) or decreased (repressive variants) mORF protein abundance. Shaded areas show distribution of p-values over 100 permutation tests; dashed line marks a one-to-one relationship. b) Common derepressive variants show a significantly increased number of GWAS hits relative to all common SNPs in 5’ UTRs (two-sided binomial test).

None of the derepressive or repressive variants we identified overlap with genes that have conserved peptide uORFs (CPuORFs) across angiosperms(44). This is possibly because the strong conservation of CPuORFs means any new mutations are subject to strong negative selection. Studies identifying CPuORFs have also been largely focused on dicots. However, many of the uORFs containing derepressive and repressive variants that we identified are conserved between members of the Andropogoneae tribe(45), although they are not significantly more or less conserved than all other uORFs (Supplemental Figure 3).

All results up to this point have been based on genes that have protein abundance data. We expanded our scope to all annotated genes in B73 and identified variants that may have repressive or derepressive function (Supplemental Data 6). Genes with derepressive variants were enriched for biological process gene ontology terms related to metabolic and biosynthetic processes (Supplemental Table 7), consistent with evidence that uORFs contribute to regulation of metabolic pathways in plants(46, 47). We identified a potential derepressive variant in the 5’ UTR of an adenosylmethionine decarboxylase, Zm00001eb184470, the ortholog of which has been shown to be regulated by uORFs in *Arabidopsis thaliana(48)*.

Beyond plants, it may be the case that certain protein families are regulated by uORFs across eukaryotes. We found derepressive variants in the uORFs of Zm00001eb252410 & Zm00001eb136490, which both encode proteins from the heat shock protein 70 family, a member of which has been shown to be regulated by uORFs in humans(49). Another derepressive variant is in the uORF of Zm00001eb070400, which is a multidrug resistance-associated protein (ABC transporter C family member 2), also shown to be regulated by uORFs in humans(50).

These results demonstrate that rare alleles have dysregulatory effects on the proteome. We have also shown that uORF translation contributes to allelic differences in mORF protein abundance between individuals. Not only do these effects show up at the protein level, but variants in uORFs are enriched for GWAS hits of metabolic and physiological traits. Because these effects relate to fundamental aspects of the translational machinery, we anticipate that these findings will extend beyond maize and be applicable in most eukaryotes. These findings can be used to engineer or predict variation in protein regulation and can potentially be applied to address problems in synthetic biology, genome editing, crop improvement, and human disease.

## Materials & Methods

### Proteomic datasets

Two distinct proteomic datasets were used in this study. The first consisted of four inbred lines (B73, Mo17, CML103, and P39) for which genome assemblies are publicly available. These were used for analyses in which accurate genomic sequence or biological replication was needed. The second dataset was generated by Jiang and colleagues(2), which consists of 98 diverse inbred lines. Protein abundance data for these lines was determined by using B73 as the reference for aligning peptides to the proteome. These data were used for analyses requiring a larger sample size and SNP sets called against a common reference genome.

### Replicated proteomics on four diverse inbreds

The plants were grown using 3:1 Metromix 900 (SunGro)/Turface MVP (Profile Products) in the greenhouse under a 16-hr light/8-hr dark photoperiod and 27 °C/ 22 °C day/ night temperatures. Third and fourth leaves of two weeks old plants were collected separately, flash frozen and stored at -80 °C. Sample preparation and mass spectrometry analysis was performed as described by McLoughlin and collaborators(51). Each genotype was analyzed with five biological and three technical replicates. Raw mass spectrometry data were analyzed against the B73 v5 proteome(52) with Proteome Discoverer (version 2.0.0.802; Thermo Fisher Scientific). Peptides were assigned by SEQUEST HT, allowing 1 missed tryptic cleavage, a precursor mass tolerance of 10 ppm and a fragment mass tolerance of 0.02 Da. Carbamidomethylation of cysteines and oxidation of methionine were used as static and dynamic modifications, respectively. Only peptides with FDRs of 0.01 (high confidence) were used for data analysis.

### Heritability calculations

We fit each peptide with the model *y*_*ij*_ = *g*_*i*_ + *e*_*ij*_ where *y*_*ij*_ was the protein abundance of peptide *i* in inbred *j, g*_*i*_ is the genetic effect of inbred *i*, and *e*_*ij*_ is the residual for peptide *i* in inbred *j*. Both terms *g* and *e* were random effects, with variances 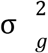 and 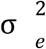 respectively. Heritability was calculated as 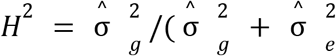.

### Protein abundance prediction

We attempted to predict peptide abundance, using several sequence-based models that predict protein or mRNA abundance. The first model we tested was described by Washburn and Wang(36), who developed it to predict which of two orthologous genes in *Zea mays* and *Sorghum bicolor* had higher mRNA expression levels. We used the same inputs, namely 1500bp of DNA sequence from each of the promoter and terminator regions of each gene, and attempted to predict which of two inbred lines had higher peptide abundance.

The second model we tested was published by Cuperus and colleagues(12), who trained their model on a library of random *Saccharomyces cerevisiae* 5’ UTRs and corresponding protein abundances. We used their pretrained model to try to predict peptide abundance in maize. Because the peptide abundances in our dataset cannot be compared between genes, we instead compared the log10-ratio of observed peptide abundances to the log10-ratio of predicted peptide abundances for each pairwise comparison between inbreds at each peptide in our dataset.

The third model we tested was based on the observation that codon bias can be predictive of protein abundance(53–55). We trained a multiple linear regression model to predict the log-ratio of peptide abundance between two inbreds, using as explanatory variables the differences in codon counts between alleles of genes encoding the same peptide.

### Uplifting Jiang proteomic data

The proteomic data generated by Jiang and colleagues(2) was uplifted to version 5 of the B73 proteome(52). The Uniprot IDs in the published data were used to obtain protein sequences, which were translated to v5 using blastp(56). Proteins with a match in the v5 proteome that had >90% identity and >90% coverage (2523 out of 2750) were kept. Similar to the four-inbred experiment described above, the protein abundances reported by Jiang and colleagues were normalized by calculating the log10-ratio against B73, in order to facilitate comparisons between genes.

### Calling HapMap SNPs on Jiang inbreds

The maize HapMap3.2.1(32) SNP data were uplifted to B73 v5 coordinates using CrossMap(57). Whole genome sequencing of 95 maize inbreds(2) was used to call SNPs at the same v5 positions and using the same methods as HapMap3.2.1(32).

### Effects of rare variants on protein abundance

To study the relationship between rare, putatively deleterious variants and protein abundance, we classified SNPs as rare if they had a minor allele frequency (MAF) < 0.02 in the maize HapMap 3.2.1 panel, which contains over 1,200 diverse maize varieties and wild relatives, and represents an independent dataset from the Jiang inbred lines for defining MAF(32).

Variants were categorized based on overlap with five annotated features: promoters (2,000bp upstream of the TSS), 5’ UTRs, 3’ UTRs, CDS, and introns. SNPs were categorized using the GenomicFeatures (58) package in R (59), based on the Zm00001eb.1 annotation of B73(52). Each combination of gene, feature, and inbred was given a binary classification for whether or not it contained a rare (MAF < 0.02) SNP allele. Within each feature, the relative protein abundance of all gene/inbred combinations that did contain a rare allele was compared to a null consisting of protein abundances in gene/inbred combinations that did not have rare variants in any of the five features. Distributions of protein abundance between the rare-variant and null categories were statistically compared by the Kolmogorov–Smirnov test(60, 61).

### Identifying translated upstream open reading frames

Two biological samples of riboseq data generated by Lei and colleagues(37) on non-stressed B73 seedlings were obtained from NCBI SRA accessions SRX845439 and SRX845455. Riboseq data consist of sequenced fragments of mRNA which are bound by ribosomes and therefore assumed to be undergoing translation. Cutadapt(62) version 1.18 was run using the parameters ‘-a CTGTAGGCACCATCAAT -m 20’ to remove adapters and discard any reads shorter than 20bp. Reads were mapped against version 5 of the B73 genome using hisat2(63) version 2.2.1 with default parameters except for ‘--trim5 1’ to remove the first bp from each read, which the original authors describe as frequently representing an untemplated addition during reverse transcription(37). The alignments were sorted and indexed with samtools(64) version 1.11, and reads that mapped to more than one location in the genome were discarded.

uORFs were computationally identified using the R(59) package ORFik(65). uORFs were identified using the pattern (ATG|TTG|CTG-3n-TAA|TAG|TGA), since translated uORFs can initiate on non-canonical start codons(66–68). Any uORFs that overlapped with annotated coding sequences were discarded, and the remaining uORFs were used as targets for read counting.

Reads overlapping computationally identified uORFs were counted by htseq(69) version 0.11.3 using htseq-count with the argument ‘--nonunique all’ so that reads were not discarded if they mapped to multiple overlapping uORFs or uORFs on different annotated transcripts. The log(FPKM + 1) values for the two biological samples were correlated, r=0.99 for CDS, r=0.75 for uORFs. The lower correlation for uORFs was primarily due to uORFs with reads in one sample but not the other; discounting those, the correlation of log(FPKM) between samples was r=0.92. These correlations were high enough that read counts from both samples were pooled and FPKM calculated on the pooled counts.

Translated uORFs were defined as computationally identified uORFs that had FPKM > 20, FPKM < 1759, total length > 15bp, and total length <333bp. These cutoffs were chosen based on the 5th and 95th percentiles of the distributions of FPKM and length across all uORFs, in order to exclude uORFs that may be in misannotated 5’ UTRs or have an unusually high number of reads mapping to them. We were relatively lenient with calling translated uORFs because, as described below, we were more interested in identifying uORFs that may be translated than obtaining highly accurate quantification of translation.

### Identifying variants that strengthen or weaken uORF start sequence

An empirical maize Kozak sequence was determined by creating a position weight matrix of the sequence spanning -3 to +4 nucleotides relative to the translation start site of all annotated maize gene models. Variants that weaken the Kozak of existing uORFs were identified by searching for variants that fell within the -3 to +4 range around the start of uORFs that show evidence of translation (described above). Two versions of the 7bp sequence around the start codon were created, one with each SNP allele, and they were scored for their similarity to the Kozak sequence using the R package Biostrings(70). Variants that decreased the similarity to the Kozak sequence by 0.5 or more were classified as ‘derepressive’.

Identification of ‘repressive’ variants was performed similarly, but with rare variants that were in the 5’ UTR of genes with no annotated uORFs, or of genes with all annotated uORFs having FPKM < 5. Because an annotated uORF was not present in all instances, unlike in the preceding paragraph it was not obvious where to position the variants within the Kozak sequence. We calculated the similarity score against the Kozak with the variant site in all positions from -3 to +4, and chose the position with the highest score, reasoning that it represented the context that was closest to making a uORF start. We then substituted the alternate allele at the SNP site and compared the Kozak score, classifying variants that increased the score by > 0.25 as ‘repressive’.

### Identifying and testing adaptive variants

Repressive and derepressive variants with MAF > 0.1 were identified by the same criteria as described above for rare alleles. One-sided Mann-Whitney tests were performed to test for increase or decrease of protein abundance associated with derepressive or repressive alleles, respectively. The results from these tests were compared to p-values from shuffling the genotypes and performing the same test 100 times for each variant.

### GWAS hit enrichment

To determine the number of physiological and metabolic traits associated with SNPs within 5’ UTR regions, a graph database was used to store and query GWAS results with additional biological information in maize. Neo4j (v4.2.0) and Cypher (v4.2.0) were used as the graph database management system and querying language, respectively(71). Results were obtained by interfacing with the database using the Neo4j diver, neo4r(59, 72). In total, 3,874 metabolite traits collected from the Goodman diversity panel(73) and 333 physiological traits collected from both the Goodman diversity and NAM panels(74, 75) were used for association testing and SNP-trait relation queries (Khaipho-Burch et al., in prep). Significant counts were obtained by filtering associations between SNP and trait with p-value < 10e-5.

### Conservation among Andropogoneae

Alignments between maize and five other grasses from the Andropogoneae tribe(45) were filtered to regions that overlap the annotated B73 5’ UTRs. uORFs, identified above, were classified based on whether at least one other species had alignments covering 90% or more of the uORF. uORFs with rare or common repressive or derepressive variants were tested against all uORFs by two-sided chi-square test to see whether repressive and derepressive variants are associated with greater or lesser conservation among the Andropogoneae.

### Gene ontology enrichment

All repressive or derepressive variants and their cognate genes were identified genomewide using the criteria previously described. Gene ontology enrichment was performed using the R package topGO(76). Fisher’s test was used to compare Biological Process GO categories for genes with repressive or derepressive variants against all genes, using the GO annotations available at https://download.maizegdb.org/Zm-B73-REFERENCE-NAM-5.0/Zm-B73-REFERENCE-NAM-5.0_Zm00001eb.1.interproscan.tsv.gz.

## Data Availability

SNP data can be found at [DOI to be made public on publication]. The mass spectrometry proteomics data for four maize inbreds have been deposited to the ProteomeXchange Consortium via the PRIDE(77) partner repository with the dataset identifier PXD026378 (Private; to be made public on publication). All other data can be found in Supplemental materials, which will be made available on publication.

## Code Availability

Code for all analyses can be found at https://bitbucket.org/bucklerlab/p_protein_diverse_maize/ (to be made public on publication).

## Acknowledgements

This material is based upon work supported by the NSF Postdoctoral Research Fellowship in Biology under Grant No. IOS-1906619, NSF Grant No. IOS-1840687, NSF Grant No. IOS-1822330, and the USDA-ARS.

## Supplemental Material

**Supplemental Figure 1:**
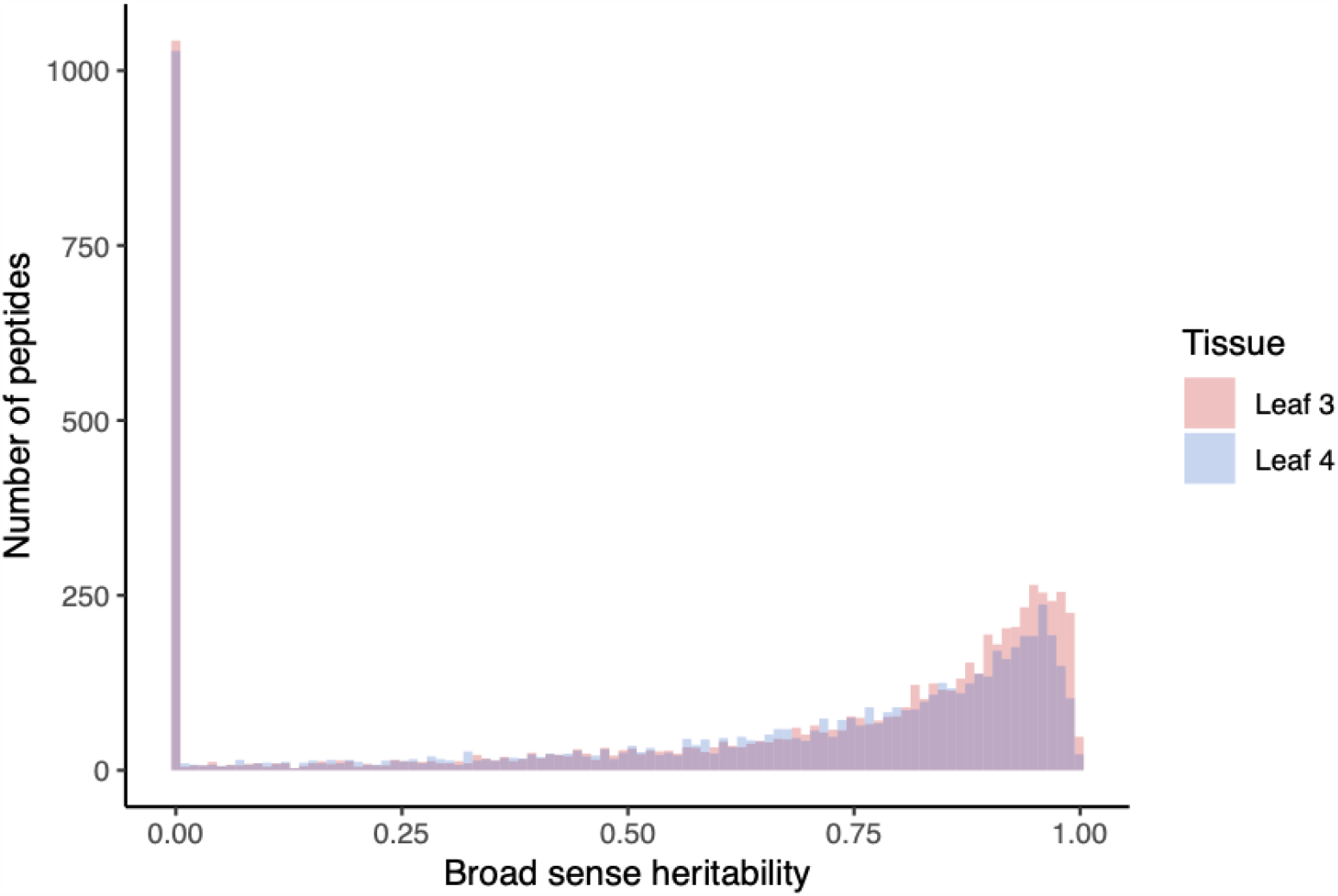
figs/supplemental/peptide_heritability.pdf. Broad-sense heritability estimates for peptides measured in four diverse inbred maize lines. Many peptides with zero heritability appear to be constitutively expressed at a constant level across individuals (ie, no genetic variance).

**Supplemental Figure 2:**
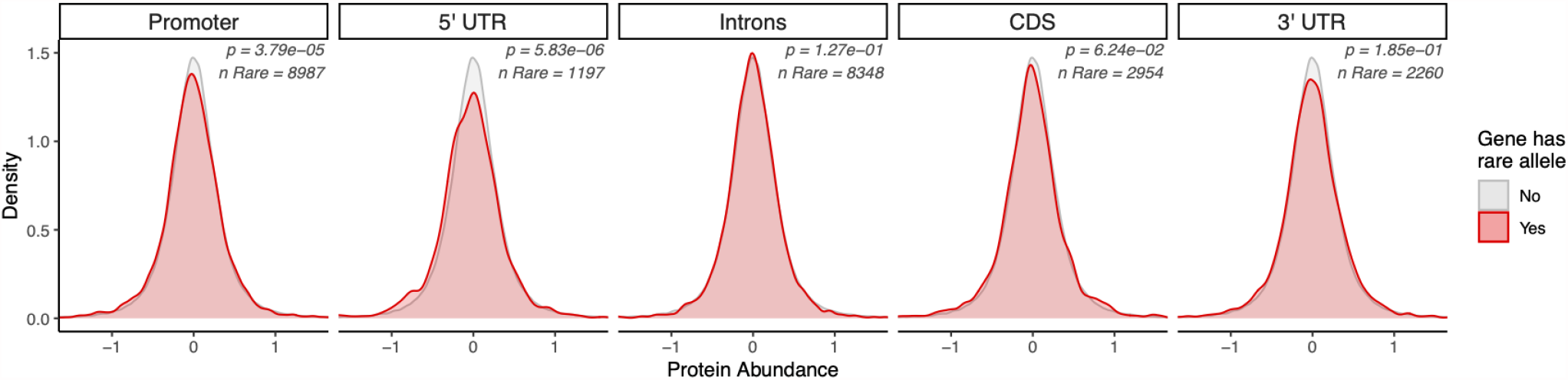
figs/supplemental/rare_alleles_genic_features_distributions.pdf. Distribution of protein abundance for individuals with rare alleles in each given region of a gene (red), compared to individuals with no rare alleles in a gene (gray). A two sided Kolmogorov-Smirnov test for differences between two distributions was used to estimate significance.

**Supplemental Figure 3:**
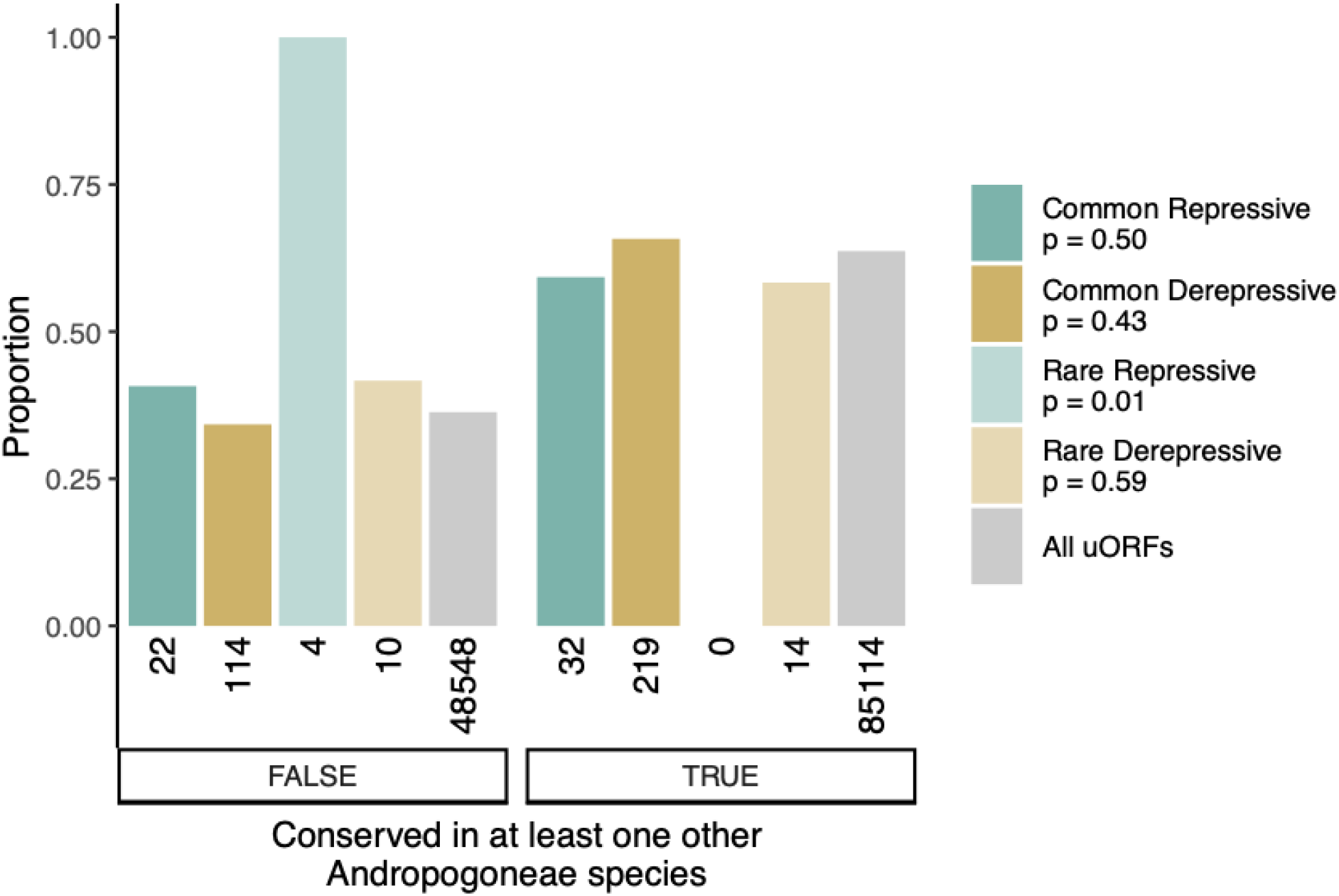
figs/supplemental/conserved_uORFs.pdf. Conservation of uORFs within five additional species from the Andropogoneae tribe. Numbers below bars are the quantity of uORFs in each category. The number of derepressive uORFs in this analysis is greater than the total number of derepressive uORF variants because for this analysis variants can be within more than one uORF. The number of repressive uORFs in this figure is lower than the total number of repressive uORF variants because not all variants were within annotated uORFs and therefore could not be scored for sequence conservation.

